# Infertility and fecundity loss of *Wolbachia*-infected *Aedes aegypti* hatched from quiescent eggs is expected to alter invasion dynamics

**DOI:** 10.1101/2020.11.25.397240

**Authors:** Meng-Jia Lau, Perran A. Ross, Ary A. Hoffmann

**Affiliations:** Pest and Environmental Adaptation Research Group, Bio21 Institute and the School of BioSciences, The University of Melbourne, Parkville, Victoria, Australia

**Keywords:** *Aedes aegypti*, *Wolbachia*, egg storage, temperature, population invasion

## Abstract

The endosymbiotic bacterium *Wolbachia* shows viral blockage in its mosquito host, leading to its use in arboviral disease control. Releases with *Wolbachia* strains *w*Mel and *w*AlbB infecting *Aedes aegypti* have taken place in several countries. Mosquito egg survival is a key factor influencing population persistence and this trait is also important when eggs are stored prior to releases. We therefore tested the viability of mosquitoes derived from *Wolbachia w*Mel and *w*AlbB-infected as well as uninfected eggs after long-term storage under diurnal temperature cycles of 11-19°C and 22-30°C. Eggs stored at 11-19°C had higher hatch proportions than those stored at 22-30°C. Adult *Wolbachia* density declined when they emerged from eggs stored for longer, which was associated with incomplete cytoplasmic incompatibility (CI) when *w*Mel-infected males were crossed with uninfected females. *w*AlbB-infected males continued to show complete CI. Females from stored eggs at both temperatures continued to show perfect maternal transmission of *Wolbachia*, but storage reduced the fecundity of both *w*Mel and *w*AlbB-infected females relative to uninfected mosquitoes. Furthermore, we found a very strong negative impact of the *w*AlbB infection on the fertility of females stored at 22-30°C, with almost 80% of females hatching after 11 weeks of storage being infertile. Our findings provide guidance for storing *Wolbachia*-infected *Ae. aegypti* eggs to ensure high fitness adult mosquitoes for release. Importantly, they also highlight the likely impact of egg quiescence on the population dynamics of *Wolbachia*-infected populations in the field, and the potential for *Wolbachia* to suppress mosquito populations through cumulative fitness costs across warm and dry periods, with expected effects on dengue transmission.

## Introduction

*Aedes aegypti* is a major vector of dengue, Zika, chikungunya and yellow fever viruses and is distributed mainly in tropical and subtropical areas [1]. *Aedes aegypti* abundance is determined by seasonal fluctuations in temperature, humidity and precipitation [2]; temperature influences mortality and development rates while rainfall affects the availability of larval habitats and egg hatching success. Long term survival of eggs is the main factor influencing population persistence especially in areas where there is a cold, dry season [3, 4]. Under ideal laboratory conditions, *Ae. aegypti* eggs can be stored for ten weeks under 27°C with more than 80% of eggs hatching when submerged [5]. In nature when eggs are exposed to harsh environments as well as predators and parasites [6], egg survival can still be around 50% after overwintering [7, 8], but high temperature and low humidity conditions increase egg mortality [9].

*Wolbachia*, a common endosymbiont in arthropods, has antiviral effects on its host and was artificially introduced into *Ae. aegypti* to help suppress arbovirus transmission [10–12]. *Wolbachia*-infected populations have been established in the field including the *w*Mel [13–15] and *w*AlbB strains [16]. This approach relies on the ability of *Wolbachia* infections to invade through cytoplasmic incompatibility (CI), when infected males cause the death of offspring when mated with uninfected females [17]. However, the *Wolbachia* infection can also induce fitness costs on its host to reduce the host population [18], while environmental conditions can influence infection dynamics in populations by affecting *Wolbachia* density, leading to other impacts like CI leakage [19] and maternal transmission failure [20]. Both these effects can increase the uninfected proportion of mosquitoes in a population across generations. A reduction in *Wolbachia* frequency in a population will reduce efficiency whereby *Wolbachia* invades a population [21–23] with consequences for successful disease control.

Given that *Wolbachia* can play an important role in arboviral disease control [13, 16], it is important to determine environmental conditions when invasion dynamics are more likely to favor *Wolbachia* to persist at a high level. In areas where *Ae. aegypti* occurs, factors such as temperature and rainfall variability will impact population abundance, through influencing egg quiescence [15, 24, 25]. Egg quiescence can be affected by high temperatures and *Wolbachia* infection status under laboratory conditions. The effect of *w*MelPop on *Ae. aegypti* eggs is particularly strong [26, 27], whereas *w*Mel- and *w*AlbB-infected eggs can survive much longer [28, 29], although fitness costs under some conditions are apparent. For example, *w*Mel-infected eggs are more sensitive than uninfected eggs to high temperatures, with lower hatch proportions when eggs are stored at 26-36 °C [19]. The impacts of rainfall variability, which is an important factor in the seasonal outbreak of mosquitoes [30], has received less attention. Moreover, environmental and *Wolbachia* effects on egg quiescence have so far been characterized within the egg life stage, whereas *Wolbachia* invasion depends on effects on host phenotypes across life stages.

In this study, we tested the viability of *w*Mel-infected, *w*AlbB-infected and uninfected eggs after storage under cold (11-19°C) and warm (22-30°C) cycling temperatures for different durations. We then tested *Wolbachia* infection density and fitness of the adults emerging from these eggs as well as on maternal transmission and CI. These experiments led to the surprising finding that many *w*AlbB-infected females showed sterility when emerging from warm-stored eggs, and females with either infection showed a reduction in fecundity. Such effects could influence the local population dynamics of *Wolbachia*, and the seasonal size of outbreaks of *Ae. aegypti* populations in areas once *Wolbachia* has established, with secondary effects on disease transmission. Our results also provide guidance on efficient accumulation of *Wolbachia*-infected eggs prior to release which can help release programs [16, 31] and they point to novel options for suppressing mosquito populations by *Wolbachia* strains in some contexts.

## Materials and Methods

### Mosquito strains and maintenance

Uninfected, *w*Mel-infected and *w*AlbB-infected *Ae. aegypti* were used in this work. The *w*Mel-infected and uninfected populations were derived from eggs collected from Cairns, Queensland, Australia [29] while the *w*AlbB-infected population was generated by Xi et al [32] and then crossed to an Australian background [29]. Mosquitoes were maintained at 26 ± 1 °C in a controlled temperature room [33] until experiments commenced. *Wolbachia*-infected populations were backcrossed to an uninfected population at least three times a year to maintain a similar genetic background in all the stocks [26]. Another three generations of backcrossing were undertaken directly before this experiment. Female mosquitoes were blood fed by a volunteer as described by the University of Melbourne Human Ethics Committee (approval 0723847). After a week, egg batches were collected on sandpaper strips, rolled up in paper towels and sealed in a plastic bag for three days before the viability of eggs was tested.

### Egg viability across storage time

We cut egg batches into small 1 cm × 3 cm pieces. Each piece contained 50 to 150 eggs which were hatched to calculate the initial hatch rate (week 0) or stored egg hatch rate under cycling temperatures of 11-19 ± 1 °C or 22-30 ± 1 °C in two incubators with a 12:12 h photoperiod. The eggs were labelled and sealed in a plastic box with a cup of saturated sodium chloride solution to balance the humidity at approximately 75% [34]. Temperature and humidity in the box were checked by data loggers (Thermochron; 1-Wire, iButton.com, Dallas Semiconductors, Sunnyvale, CA, USA) (Supplementary Figure 1). After storage for 0, 1, 4, 8, 11, 14 or 16 weeks, four to six batches of eggs per population and temperature cycle were submerged in reverse osmosis (RO) water with TetraMin^®^ fish food tablets (Tetra, Melle, Germany) *ad libitum* and yeast (to stimulate the hatching) and a second immersion was carried out with a one-day drying interval to hatch any viable eggs that had failed to hatch in the first immersion [35]. Intact and hatched eggs (with a detached egg cap) were counted to indicate dead or surviving larvae. After hatching, larvae were reared until the adult stage for further testing.

### *Wolbachia* density across storage time

*Wolbachia*-infected eggs before (0 weeks) and after cycling temperature treatments (8, 11, 14, or 16 weeks) were hatched. Two-day-old larvae were density controlled to a maximum of 30 larvae in trays of 300 mL RO water in each treatment and reared to adulthood. Eleven 2-day-old females per treatment were stored in 100 % ethanol for *Wolbachia* screening. Individual DNA was extracted in 200 μL 5% Chelex 100 Resin (Bio-Rad Laboratories, Hercules, CA) [36], and diluted ten times for quantitative PCR analyses. Mosquito-specific (*mRpS6*) primers, *Ae. aegypti*-specific (*aRpS6*) primers and *Wolbachia*-specific primers (*w1* primers for *w*Mel-infected or *wAlbB* primers for *w*AlbB-infected) were used in the same run for each sample [29, 37, 38]. Three technical replicates were completed for each sample with two requirements: first, the difference of the crossing point (ΔCp) between *mRpS6* and *aRpS6* primers, which was used to indicate the quality of each run, had to be less than 1; and second, the variance of ΔCp between the three runs was expected to be less than 0.1. Relative density was calculated by power 2 transformation of ΔCp differences between *aRpS6* and *Wolbachia* primers and technical replicates were averaged before testing treatment differences (storage time, *Wolbachia* infection type, temperature).

### Cytoplasmic incompatibility

*w*Mel- and *w*AlbB-infected eggs that not been stored (preserved under constant 26 °C for less than three weeks, regarded as untreated) or had been stored (treated) for 14 weeks at 11-19 °C and 10 (*w*AlbB) or 11 (*w*Mel) weeks at 22-30 °C were hatched and larvae were reared to adulthood for crossing experiments. Wing lengths were measured for seven to ten randomly selected females from each treatment as a measure of mosquito size. Eight replicate cages, each consisting of five uninfected untreated females crossed with five *Wolbachia*-infected treated males, were set up to test cytoplasmic incompatibility (CI), while four replicates involving crosses with *Wolbachia*-infected untreated males were set up as controls. Eight replicates, each with five *Wolbachia*-infected treated females crossed with five *Wolbachia*-infected untreated males, were set up to test for CI rescue, while four replicates of *Wolbachia*-infected untreated females crossed with untreated males were set up as controls. After adults emerged, they were left with a 10% sucrose cup and water cup for four or five days to ensure that females had matured and mated. After blood feeding, females were allowed to oviposit on Norton®Master Painters P80 sandpaper (3.8 × 18 cm; Saint-Gobain Abrasives Pty. Ltd., Thomastown, Victoria, Australia) which was half submerged into water which had been used to rear larvae. These egg strips were collected at the fifth and seventh day and hatched on the ninth day after blood feeding.

### Maternal transmission

To test for maternal transmission, 20 *Wolbachia*-infected female mosquitoes that had been stored for 14 weeks at 11-19 °C and 10 (*w*AlbB) or 11 (*w*Mel) weeks at 22-30 °C were crossed to uninfected males that had not been stored before blood feeding. Females were individually isolated in a 70 mL cup after blood feeding and allowed to lay eggs for one week. After laying eggs, mothers were stored in ethanol for *Wolbachia* screening. Mothers who died before being stored were excluded from the experiment. We selected six to seven mothers per treatment which each had more than 30 offspring alive, and we screened the mother and eight to ten offspring for *Wolbachia* infection. *w*AlbB-infected females from the 22-30 °C treatment that failed to lay any eggs were dissected after being stored in ethanol to look for the presence of mature eggs. Since no *w*AlbB-infected females stored for 10 weeks at 22-30 °C laid eggs in this experiment, we instead tested maternal transmission from *w*AlbB-infected females stored for 11 weeks at 22-30 °C in a second experiment (see below).

### Female fertility

We further investigated the loss of fertility in *Wolbachia*-infected females derived from warm (22-30 °C) stored eggs by hatching batches of *w*AlbB-infected (stored for 0, 6, 9 or 11 weeks), *w*Mel-infected (0, 9 and 11 weeks) and uninfected eggs (0, 6, 9 and 11 weeks). To understand the role males played, females hatched at each time point were (1) crossed to males from the same treatment, (2) crossed to males with the same *Wolbachia* infection type that were not stored, and (3) crossed to uninfected untreated males. Between 30-42 females from each cross were isolated in a 70 mL cup and provided with a strip of sandpaper to lay eggs. Eggs were collected once at the sixth day and hatched at the ninth day after blood feeding to determine fecundity and egg hatch proportions. Females that died before egg collection were excluded from analyses.

We determined the prevalence of female infertility by scoring the proportion of females in each cross that did not lay eggs on the collection day. We dissected female mosquitoes that failed to produce eggs from each cross at 9 weeks, and at 11 weeks we randomly selected and dissected up to ten female mosquitoes from each cross. We identified female insemination status by looking at their spermathecae under a compound light microscope (Motic B1 series, Australian Instrument Services Pty. Ltd., Australia). Each female has three spermathecae, one larger one and two smaller ones. The spermathecae containing sperm were distinguished from those that did not based on their texture and transparency, with spermathecae containing sperm having a dark and solid appearance with visible sperm swirling around the perimeter (Supplementary Figure 2, Supplementary video 1).

### Statistical analysis

All data analysis and visualization were undertaken with R v.3.6.0. We performed Kaplan-Meier survival tests and log rank tests [39] to compare treatment effects on changes in egg viability across time, and used the Benjamini p-value adjustment method for pairwise comparisons. We used ANOVAs to compare treatment effects for normally distributed data including fecundity. We log transformed *Wolbachia* density and used logistic regression to analyze treatment and storage time effects on proportional data such as egg hatch and female infertility. We ran Kruskal-Wallis rank tests to analyze treatment effects of hatch proportions in the maternal transmission experiment. Hatch proportions were arcsine square root transformed before computing 95% confidence intervals but these were then back transformed before being presented (Supplementary Table 3).

## Results

### Egg viability across storage time

We compared the viability of uninfected, *w*Mel-infected and *w*AlbB-infected eggs when stored at 11-19°C or 22-30°C for up to 16 weeks. There was a significant difference of infection treatment (log rank test, χ^2^ = 310, df = 2, p < 0.001) when eggs were stored at 22-30°C (Figure 1), with the *w*AlbB-infected eggs dying faster than the uninfected and *w*Mel-infected eggs (pairwise comparisons, *w*AlbB-infected : uninfected, p < 0.001; *w*AlbB-infected : *w*Mel-infected, p < 0.001; *w*Mel-infected : uninfected, p = 0.004). Eggs were viable for longer under 11-19 °C (all colonies, p < 0.001), and there were only marginally significant differences between infection treatments at this temperature (log rank test, χ^2^ = 6.4, df = 2, p = 0.04); *w*Mel-infected eggs died faster than uninfected eggs (pairwise comparisons, *w*AlbB-infected : uninfected, p = 0.287; *w*AlbB-infected : *w*Mel-infected, p = 0.626; *w*Mel-infected : uninfected, p = 0.027).

**Figure 1.**
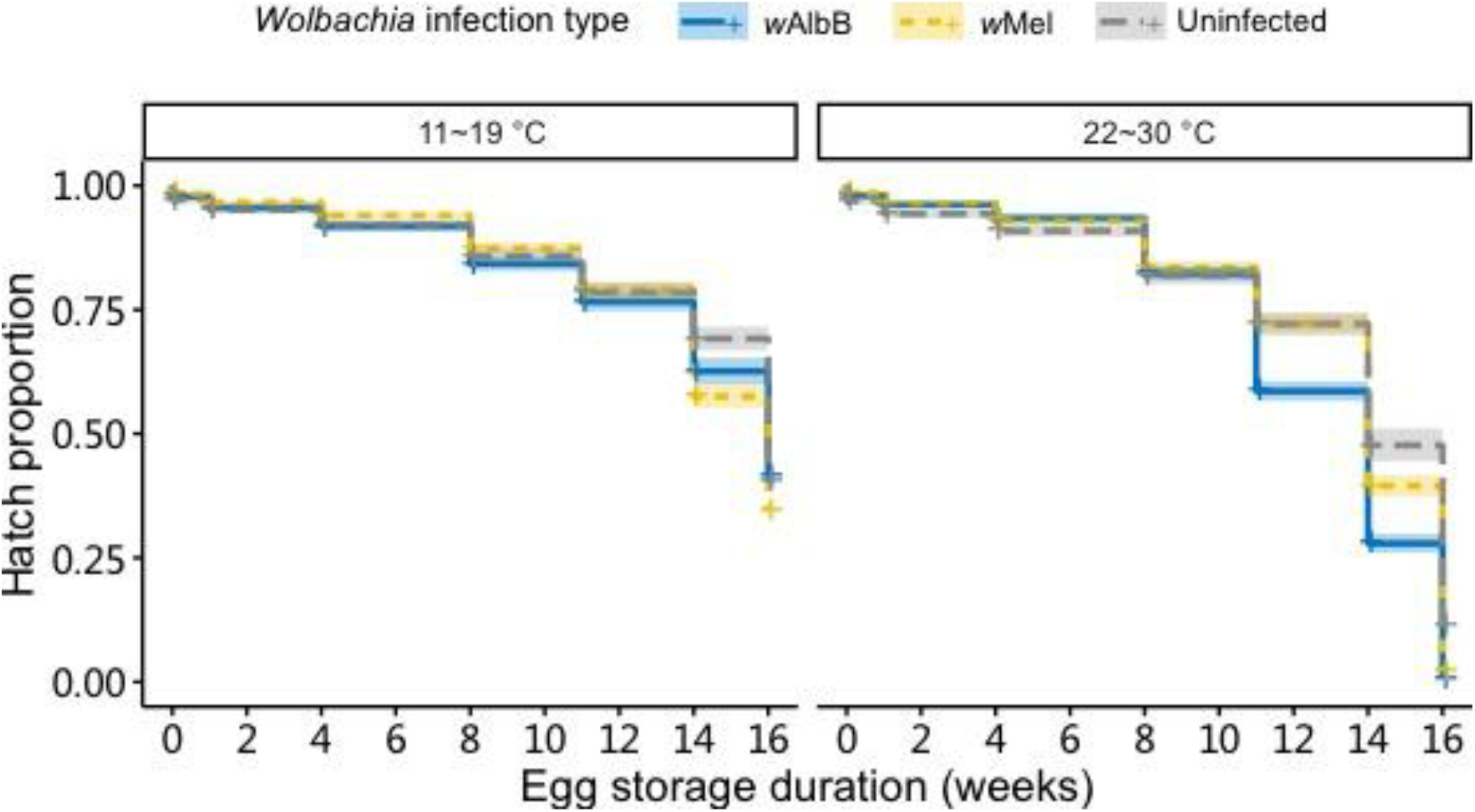
Survival of eggs based on hatch proportions of uninfected, *w*AlbB-infected and *w*Mel-infected *Aedes aegypti* eggs stored under cycling temperatures of 11-19 °C or 22-30 °C for 0, 1, 4, 8, 11, 14 or 16 weeks. Shaded areas represent 95% confidence intervals.

### *Wolbachia* density across storage time

We found that *Wolbachia* density (log transformed) in female adults hatching from quiescent eggs was significantly affected by storage time and *Wolbachia* infection type, but not by temperature (three-way ANOVA, storage time: F_1,176_ = 26.869, p < 0.001; *Wolbachia* infection type: F_1,176_ = 10.363, p = 0.002; temperature: F_1,176_ = 0.618, p = 0.433). Density tended to decrease with egg storage time (Figure 2). There was a significant interaction among the three variables (three-way interaction, F_1,176_ = 8.609, p = 0.004) but two-way interactions were not significant (p > 0.05). When different mosquito populations were considered separately, both the density of *w*Mel and *w*AlbB-infected mosquitoes were related to storage time but not storage temperature (storage time: both p < 0.001; storage temperature: *w*Mel-infected: F_1,107_ < 0.001, p = 0.996; *w*AlbB-infected: F_1,69_ = 0.578, p = 0.450).

**Figure 2.**
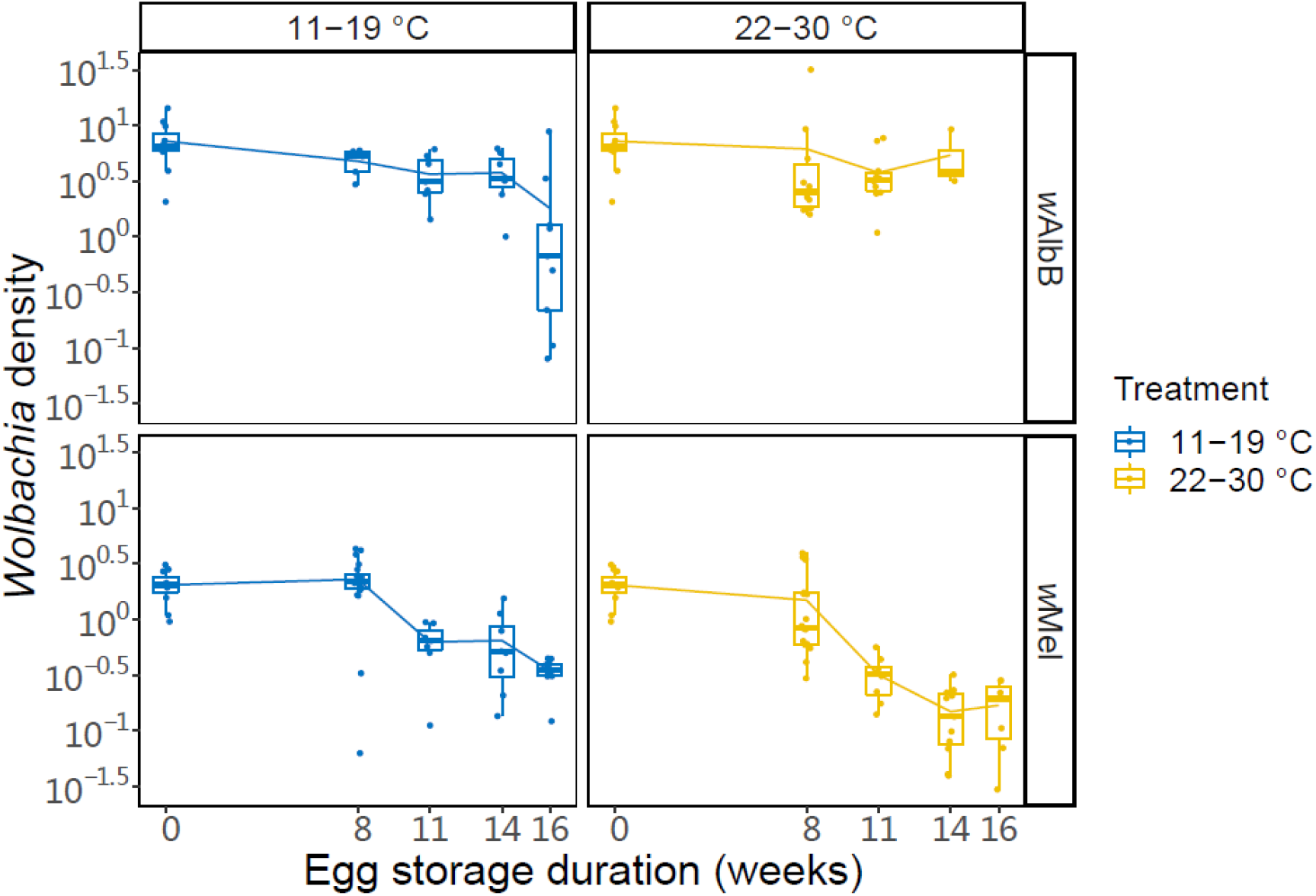
Box plots of **l**og relative *Wolbachia* density of *w*Mel-infected or *w*AlbB-infected 2-day-old female mosquitoes hatched from eggs stored under temperature cycles of 11-19°C or 22-30°C for 0, 8, 11, 14 or 16 weeks. Points represent individuals.

### Cytoplasmic incompatibility

The *w*AlbB-infected males caused complete CI with uninfected females (no eggs hatched) regardless of egg storage duration. For *w*Mel-infected *Ae. aegypti*, incomplete CI occurred when males were reared from eggs stored at either 11-19 °C or 22-30 °C. No eggs hatched from crosses where *w*Mel-infected males were hatched from unstored eggs. However, hatch proportions in CI crosses were 0.35% and 0.26% when *w*Mel-infected fathers were stored at 11-19 °C and 22-30 °C respectively (Supplementary Table 1), suggesting a minor loss of CI.

We performed crosses between *Wolbachia*-infected treated females (hatched from eggs stored at 11-19 °C or 22-30 °C) and *Wolbachia*-infected untreated males to test for CI rescue. The number of eggs laid by females was impacted by their egg storage treatment (ANOVA, F_2,34_ = 39.593, p < 0.001), but not by their *Wolbachia* infection type (F_1,34_ = 1.423, p = 0.241). *Wolbachia*-infected females derived from eggs stored under 22-30 °C for 10/11 weeks laid approximately 70% and 45% fewer eggs for *w*AlbB-infected and *w*Mel-infected respectively than when stored under 11-19 °C (Figure 3). Egg hatch proportions from these crosses were unaffected by the egg storage treatment of the female parents (Figure 3B, Logistic regression: F_2,34_ = 2.669, p = 0.084), or their *Wolbachia* infection type (F_1,34_ = 0.130, p = 0.721), suggesting that *Wolbachia*-infected females are able to restore compatibility with *Wolbachia*-infected males regardless of storage conditions.

**Figure 3.**
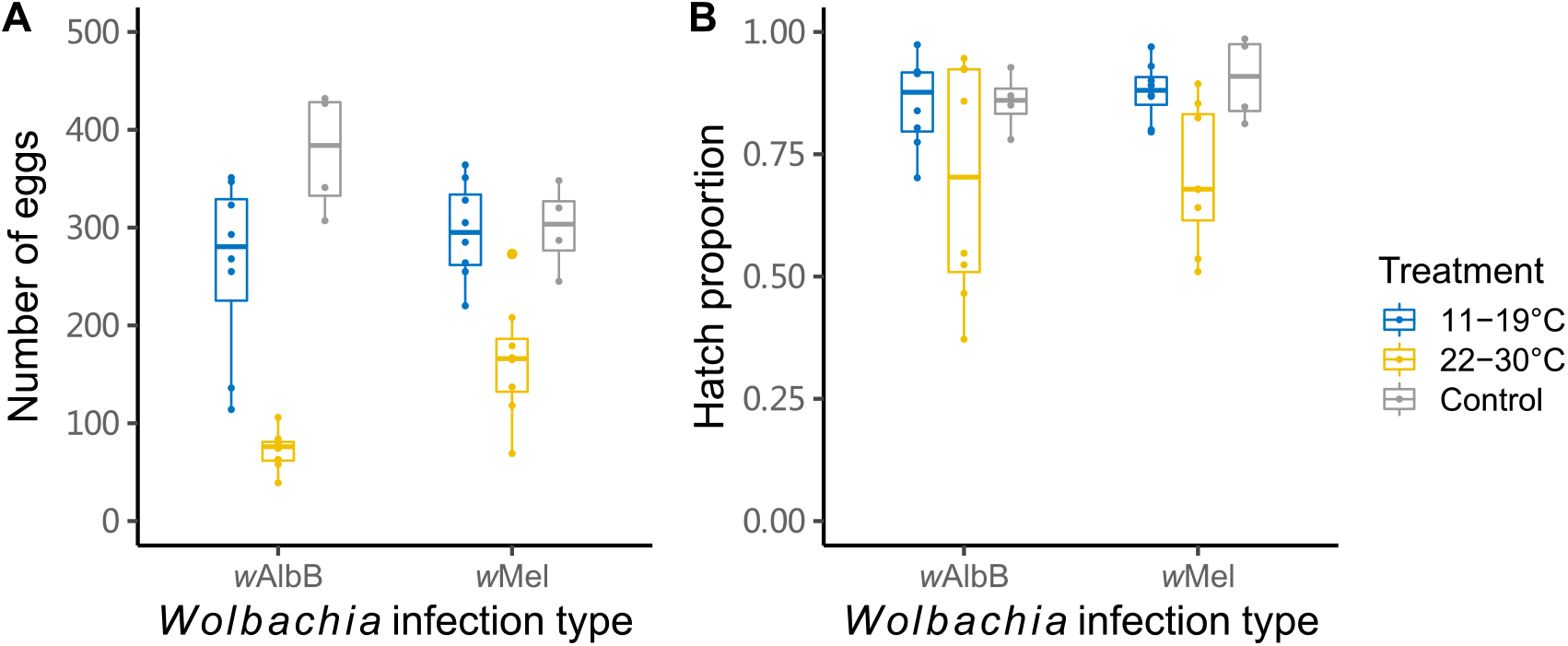
(A) Number of eggs laid by females and (B) their hatch proportions from crosses between *Wolbachia*-infected females and *Wolbachia*-infected males. *Wolbachia*-infected females were derived from eggs stored under 26°C for one week (control) or a cycling temperature of 11-19°C for 14 weeks or 22-30 °C for 10 (*w*AlbB-infected) or 11 (*w*Mel-infected) weeks. Each data point represents the total number of eggs laid and their hatch proportion from a replicate cage of five females and five males.

We measured the wing length of female mosquitoes to estimate their body size. No significant differences were found for *w*Mel or *w*AlbB between the two temperature treatments and untreated samples (ANOVA, *w*Mel: F_2,23_ = 0.578, p = 0.569; *w*AlbB: F_2,23_ = 3.153, p = 0.062). There were weakly significant differences in wing length between untreated *w*Mel-infected, *w*AlbB-infected and uninfected females (F_2,19_ = 4.62, p = 0.023), where *w*AlbB-infected females were somewhat larger than females from the other two populations (Supplementary Table 2).

### Maternal transmission

*Wolbachia*-infected females stored at 22-30 °C or 11-19 °C transmitted the infection to the next generation at a frequency of 100% regardless of *Wolbachia* infection type and egg storage temperature. In our first test of *w*AlbB-infected females stored at 22-30 °C, all twelve isolated females did not lay eggs, and no mature eggs were observed in their body upon dissection. We then counted the number of eggs laid by *w*AlbB-infected females stored at 11-19 °C and *w*Mel-infected females from both treatments and their hatch proportions (Supplementary Table 3). No significant differences of hatch proportion were found (Kruskal-Wallis rank tests, comparison between *w*Mel-infected and *w*AlbB-infected mosquitoes under 11-19 °C: χ2 = 0.776, df = 1, p = 0.379; comparison between 11-19 °C and 22-30 °C for *w*AlbB-infected mosquitoes: χ2 = 3.564, df = 1, p = 0.059). However, *w*Mel-infected females produced fewer eggs when stored at 22-30 °C compared to 11-19 °C (ANOVA, F_1,27_ = 4.875, p = 0.036).

### Female fertility

The results in the previous experiments suggest fertility effects from storage in that *Wolbachia*-infected females produced low egg numbers (or did not lay eggs at all) when stored as eggs under 22-30 °C for a period. To further investigate the apparent loss of female fertility under egg storage, we determined the proportion of *Wolbachia*-infected and uninfected females that did not lay eggs when female parents were derived from eggs stored under 22-30 °C. When *Wolbachia* infection type was considered separately, there were differences for the *w*AlbB-infected strain for both egg storage duration (F_3,6_ = 185.138, p < 0.001) and cross (F_2,6_ = 8.985, p = 0.016), whereas *w*Mel-infected and uninfected strains showed no significant effects of these factors (all p > 0.05). The proportion of *w*AlbB-infected females not laying eggs increased dramatically with egg storage duration, with over 75% of *w*AlbB-infected females not producing eggs by the final assessment time (Figure 4). All dissected females were inseminated (Supplementary Figure 2) but lacked visible eggs in their ovaries, indicating that infertility was not due to a lack of mating.

**Figure 4.**
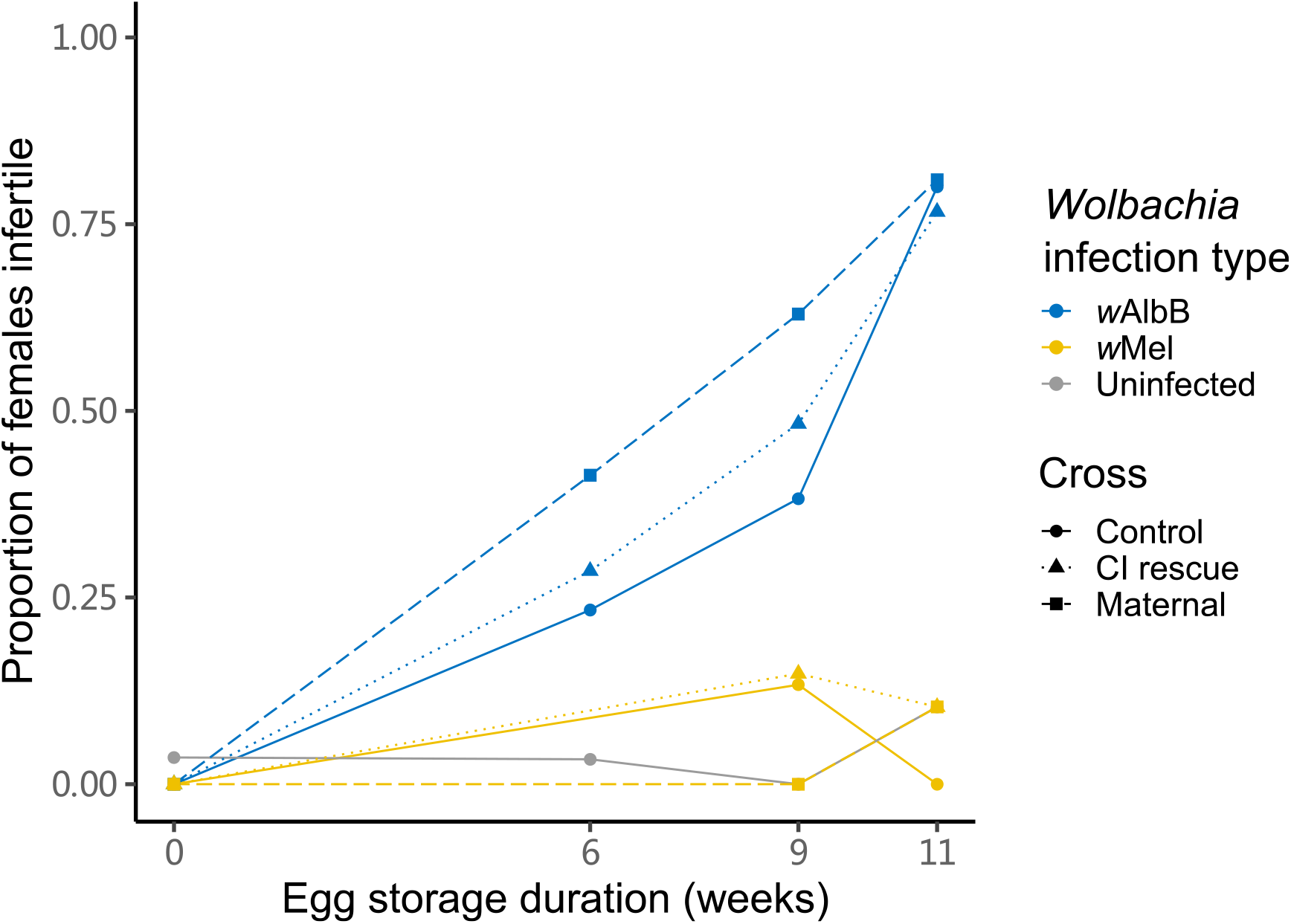
Loss of female fertility when mosquitoes were derived from eggs stored under a cycling temperature of 22-30°C for 0, 6, 9 or 11 weeks. Females hatched at each time point were crossed to males from the same treatment (Control), crossed to males with the same *Wolbachia* infection type that were not stored (CI rescue) and crossed to uninfected untreated males (Maternal). Each data point is based on the proportion of 22-42 females that did not lay eggs.

For the fertility of females that laid eggs, there were effects of *Wolbachia* infection type (ANOVA, F_2.537_ = 18.361, p < 0.001) and egg storage duration (F_1.537_ = 90.650, p < 0.001) on the number of eggs per female laid but no effect of cross (F_2.537_ = 0.038, p = 0.962). Both *w*Mel- and *w*AlbB-infected females laid fewer eggs with increasing egg storage duration in all crosses (Figure 5A). In contrast, uninfected female fecundity did not decline. For the hatch proportion of these eggs, effects were induced by *Wolbachia* infection type (Figure 5B, Logistic regression, F_2.537_ = 17.100, p < 0.001), cross (F_2.537_ = 5.734, p = 0.003) but not egg storage duration (F_1.537_ = 0.523, p = 0.470).

**Figure 5.**
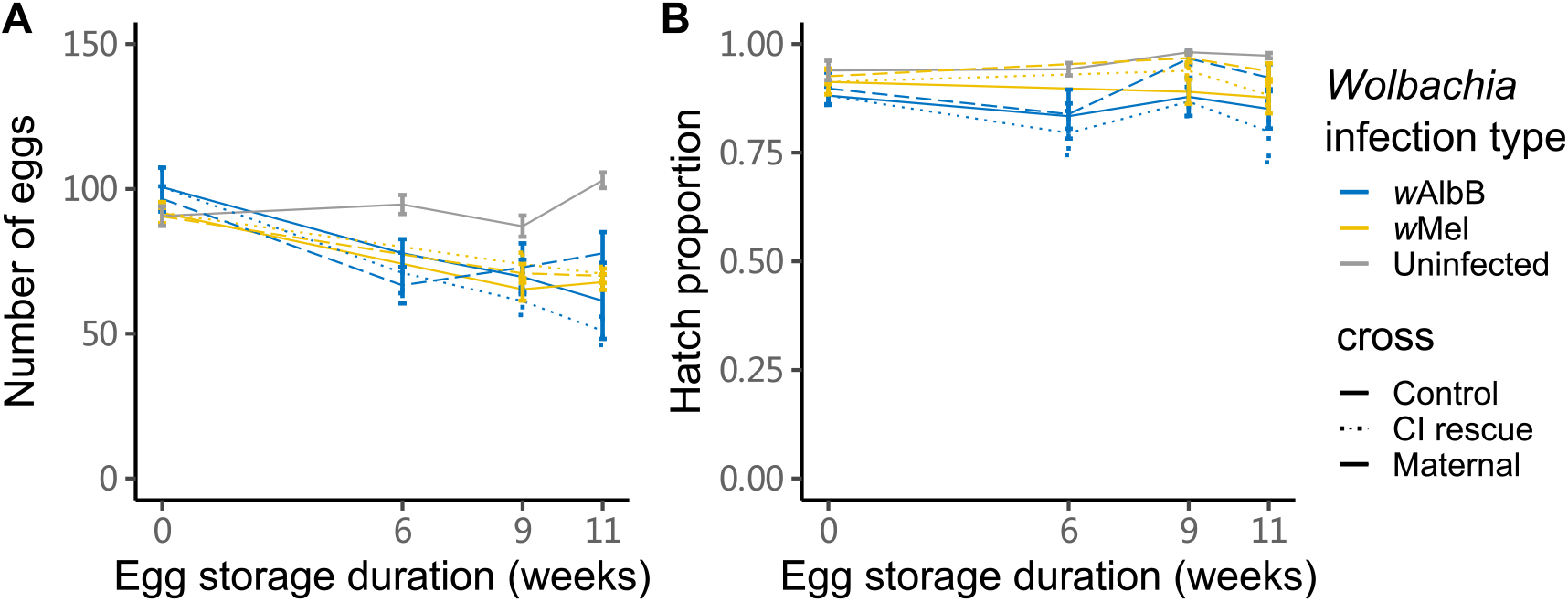
(A) Fecundity and (B) egg hatch proportions of eggs from female mosquitoes that were derived from eggs stored under a cycling temperature of 22-30°C for 0, 6, 9 or 11 weeks. Females hatched at each time point were crossed to males from the same treatment (Control), crossed to males with the same *Wolbachia* infection type that were not stored (CI rescue) and crossed to uninfected untreated males (Maternal). The data exclude females that did not lay eggs (see Figure 4). Error bars represent means ± standard errors.

## Discussion

*Aedes aegypti* mosquitoes carrying the *w*Mel or wAlbB strains of *Wolbachia* have the potential to reduce dengue transmission through decreased mosquito vector competence [16, 40], and there is already good evidence that both strains are having such impacts in *Wolbachia*-invaded release areas [13, 16]. These releases can involve adults or eggs. Releases involving eggs, where containers of eggs are hatched directly in the field, are desirable for *Wolbachia* release programs because they do not require local mass-rearing facilities [13, 16, 41] and allow eggs to be accumulated across several weeks before being transported for release. Although most releases to date have involved the *w*Mel strain, *w*AlbB was used for releases in Malaysia [16] based on its desirable characteristic of being stable in its density and phenotypic effects under high temperatures [42].

In this study, we investigated the effects of long-term storage on *Ae. aegypti* eggs infected with *Wolbachia* strains *w*Mel or *w*AlbB. Eggs survived longer under cooler environments; more than 50% of eggs from all strains hatched after storage for 14 weeks at 11-19°C while hatch proportions were below 50% when stored at 22-30°C for the same amount of time. Although the *Wolbachia* density of both *w*Mel and *w*AlbB-infected adults decreased with storage time, density was not affected by storage temperature, suggesting that density did not contribute to the decrease in egg viability under warmer temperatures. Nevertheless, we suspect that the decreased density contributed to incomplete CI for *w*Mel-infected males stored under both temperature treatments given the strong relationship between CI and *Wolbachia* density previously reported for *w*Mel [19]. Despite decreased density, the maternal transmission of *Wolbachia* remained stable after eggs had passed through a quiescent phase, regardless of storage temperature.

However, *Wolbachia* infections induced substantial costs to fertility when adults were reared from eggs stored under 22-30°C for 10 or 11 weeks. Both *w*Mel and *w*AlbB-infected females that were stored as eggs at 22-30°C produced fewer eggs. In a second experiment where we tested individual females, a high proportion of *w*AlbB-infected females suffered infertility when stored as eggs at 22-30°C, regardless of the male used in crosses. In contrast, *w*Mel-infected and uninfected females retained high rates of fertility throughout the experiment. Dissection of *w*AlbB-infected females that failed to lay eggs showed that all females were inseminated but lacked mature eggs in their abdomens despite being blood fed. Previous studies suggest that *w*AlbB has only a minor negative impact on egg viability during the first two months of storage [43]. However, these costs may be underestimated because a high proportion of females hatching from stored eggs are infertile. Furthermore, both *w*AlbB and *w*Mel-infected females laid fewer eggs when hatching from stored eggs though the hatch rate remain high. These effects appear to be unrelated to larval development, since body sizes were similar across *Wolbachia* infection types and egg storage treatments (Supplementary Table 2). *Wolbachia* infections compete with their hosts for key nutrients required for egg production [44] which may be exacerbated when female parents are stored as eggs. Egg provisioning or hormone modulation may also be impacted by *Wolbachia* under stress [43, 45], but this requires further investigation.

The survival and fertility effects generated in the quiescent *Ae. aegypti* eggs seem likely to impact the spread and maintenance of *Wolbachia* in populations in certain situations. Mosquitoes with *w*AlbB may not invade readily if releases happen in areas where larval habitats used by *Ae. aegypti* are intermittent, requiring long periods of egg quiescence. In such environments, the threshold frequency of *w*AlbB in a population that needs to be exceeded to get invasion may increase above the currently estimated point (for *w*Mel) of 20-30% [31]. Releases based on *w*AlbB eggs rather than adults (such as carried out at one site in Malaysia [16]) could also be much less effective than expected if they use eggs that have been stored too long, unless these have been stored under cool conditions. *Aedes aegypti* is an opportunistic breeder, taking advantage of locally suitable conditions which are expected to vary spatially and temporally depending on rubbish accumulation, defective building spaces and defective water and sewerage tanks [46–48]. Because of these differences, the contribution of mosquitoes from quiescent eggs to the adult population is expected to vary locally. Releases across different locations show that rates of *Wolbachia* invasion can vary for both *w*AlbB [16] and *w*Mel [46]. While various factors may be responsible for site-to-site variation in invasion rates and success including density dependent processes [21], fitness costs associated with quiescence are also likely to be important, particularly for *w*AlbB.

Once *Wolbachia* invasions have been completed, there is the potential for *Wolbachia* infections to decrease the overall size of mosquito populations due to fitness costs [27, 49]. Our data suggest that this is particularly likely in locations with strong seasonality, where quiescent eggs persisting during the dry season are more likely to contribute to *Ae. aegypti* population continuity. A strategy for crashing *Ae. aegypti* populations during the dry season based on the reduced hatch rate of infected quiescent eggs was originally proposed several years ago, based around the *w*MelPop infection which induces a particularly large fitness costs in quiescent eggs [26, 50]. However, this strain cannot easily invade mosquito populations [10, 18] whereas the *w*AlbB strain does establish successfully [16] and may carry fitness costs appropriate for such an intervention.

Given that we only tested infertility effects in females stored for up to 11 weeks under warm environment, it is unclear whether this effect will occur under cooler environments or occur in *w*Mel as well when stored for a longer period. The original host of *w*AlbB, *Aedes albopictus*, has higher cold tolerance than *Ae. aegypti* and prefers cooler temperatures [51–53]. The *w*AlbB infection may provide fitness benefits to *Ae. aegypti* under cool environments, which points to the *w*AlbB-infected *Ae. aegypti* being relatively better adapted to cool environments than uninfected mosquitoes [54]. Under hot environments, *w*AlbB is relatively more stable than *w*Mel in terms of density and its ability to induce cytoplasmic incompatibility [42]. The infertility effect we describe here was detected after only six weeks of egg storage under a milder temperature range than the hot temperatures tested previously. Nevertheless, the ability of *w*AlbB to maintain a relatively high density in quiescing eggs may result in the accumulation of damage in such eggs which leads to fitness effects at a later life stage, producing female infertility and a reduction in fecundity.

In summary, we have tested the performance of *w*Mel, *w*AlbB-infected and uninfected *Ae. aegypti* after eggs were stored under cycling temperatures of 11-19°C and 22-30°C. We found that, compared to warm environments that are normally used to culture mosquitoes [33, 55], cooler environments may be better for egg storage prior to *Wolbachia* releases. Long-term storage under warm environment greatly reduces the fertility of hatched females that carry *w*AlbB, with consequences for *Wolbachia* invasion success. Our study helps to guide conditions for appropriate storage of *Wolbachia*-infected eggs, inform release timing to coincide with periods where quiescent eggs contribute little to population dynamics, and help to explain *Wolbachia* invasion success or failure in different environments [16].

## Acknowledgements

This research was supported by the National Health and Medical Research Council (1132412, 1118640, www.nhmrc.gov.au) and the Wellcome Trust (108508, wellcome.ac.uk) for financial support. We thank Heng Lin Yeap from the Commonwealth Scientific and Industrial Research Organisation (CSIRO) and Marianne Coquilleau from the University of Melbourne for their advice and assistance.

